# Structure-guided design of a *Plasmodium vivax* Duffy binding protein-based vaccine immunogen

**DOI:** 10.1101/2024.06.23.600241

**Authors:** Natalie M. Barber, Tossapol Pholcharee, Amelia M Lias, Doris Quinkert, James Nugent, Lloyd D. W. King, Simon J. Draper, Matthew K. Higgins

## Abstract

*Plasmodium vivax* remains one of the major causative agents of human malaria and a vaccine is urgently required. It is an obligate intracellular parasites and replication within red blood cells is essential for development of disease and for transmission. The interaction between PvDBP on the parasite surface and the DARC receptor on human reticulocytes is essential for a *Plasmodium vivax* blood stage infection. Human vaccination with the RII region of PvDBP slowed parasite replication, showing that PvDBP is a promising vaccine candidate. However, it did not induce sterile protection, and further development is required to generate a vaccine which protects from clinical malaria. In this study, we develop a vaccine immunogen containing a region of PvDBP-RII, known as subdomain 3, which contains the epitope for a broadly-reactive growth-inhibitory antibody, DB9. We used structure-guided approaches to resurface subdomain 3 such that it folds as an isolated molecule. We show that this engineered subdomain 3 is more stable and more easily produced than PvDBP-RII and induces a more effective growth-inhibitory antibody response. We therefore present an improved PvDBP-based immunogen for use in blood stage vaccines to prevent malaria due to *Plasmodium vivax*.

**One sentence summary:** Structure-guided design leads to a more effective Duffy-binding protein-based vaccine immunogen to prevent *Plasmodium vivax*.

## Introduction

*Plasmodium vivax* is the predominant cause of human malaria outside Africa, leading to around 14.5 million annual cases1. While it does not receive the same attention as its more deadly relative, *Plasmodium falciparum*, it causes significant human suffering and an effective vaccine is urgently required^2,3^. The blood stage of the *Plasmodium vivax* life cycle is a promising point of intervention. The symptoms of malaria occur as the parasite invades and replicates within human reticulocytes^4^. In addition, differentiation of blood-stage parasites into gametocytes allows their uptake and development in mosquitos. A vaccine which prevents reticulocyte invasion would therefore prevent the symptoms and transmission of malaria^5^.

Reticulocyte invasion requires binding of the *Plasmodium vivax* Duffy binding protein, PvDBP, to the human Duffy antigen/receptor for chemokines, DARC, which is found on the reticulocyte surface^6^. The importance of this interaction in vivax malaria is emphasised by the effect of the Duffy-negative phenotype^7^. This polymorphism in DARC is common through much of Africa, and there is a close geographical correlation between Duffy-negativity and the reduced prevalence of *Plasmodium vivax*^2^. Indeed, knockout of the orthologue, PkDBPα, prevents invasion of Duffy-positive erythrocytes by transgenic *Plasmodium knowlesi*^8-10^. PvDBP is the most developed blood stage vaccine candidate in the quest to prevent vivax malaria.

PvDBP has a large modular ectodomain. Within this lies a ∼350 amino acid residue Duffy-binding-like domain known as PvDBP-RII, which interacts with the 60 residue DARC ectodomain^11,12^. Immunisation of mice, rabbits and non-human primates with PvDBP-RII induces inhibitory antibodies that block binding of PvDBP to DARC^13,14^. In humans, high-titres of naturally-acquired PvDBP-RII-targeting antibodies reduce DARC binding *in vitro* and are associated with decreased risk of *Plasmodium vivax* infection^15^, lower parasite densities and reduced risk of clinical malaria^16,17^. Immunisation of human volunteers with recombinant viral vectors expressing PvDBP-RII induces strain-transcending antibodies which prevent PvDBP-RII from binding to DARC, while human antibodies, from either vaccination or natural infection, inhibit invasion^9,18^. More recently, human volunteers have been vaccinated with PvDBP-RII, either delivered through a viral vector or as a protein with the Matrix M adjuvant. On challenge with *Plasmodium vivax* parasites, volunteers vaccinated with the protein vaccine showed a reduction in mean parasite multiplication rate of ∼50% when compared with unvaccinated controls^19^.

While human vaccine trials with PvDBP have shown efficacy, they also indicate that the vaccine induced responses from PvDBP-RII are not sufficient to allow sterile protection. These findings encourage a rational, structure-guided approach to the design of improved PvDBP-based immunogens. Structural studies have shown that PvDBP-RII consists of three subdomains, with subdomains 1 and 2 forming a single structural unit and subdomain 3 as a separate unit^20^. The ectodomain of DARC binds to PvDBP-RII, with the interface centred around a sulphated tyrosine residue^21^ (Figure 1a,b). In *in vitro* studies, DARC binding has been shown to induce PvDBP-RII dimer formation, with the DARC peptide located at the dimerization interface^22,23^. While there is currently no data to show that dimerization is functionally relevant *in vivo* during invasion, the binding site for DARC and the dimerization surface are both proposed as potential sites to target with vaccine-induced antibodies.Indeed, screening a linear peptide array with non-inhibitory and inhibitory human serum identified peptides which recognise antibodies found specifically in inhibitory serum^24^ and are located in regions of subdomain 2 involved in PvDBP-RII dimerisation and DARC^19-30^ binding^23^. In contrast, monoclonal antibodies derived from PvDBP-RII-immunised mice which prevent PvDBP-RII from binding to DARC *in vitro*, bind to subdomain 3^25^.

**Figure 1:**
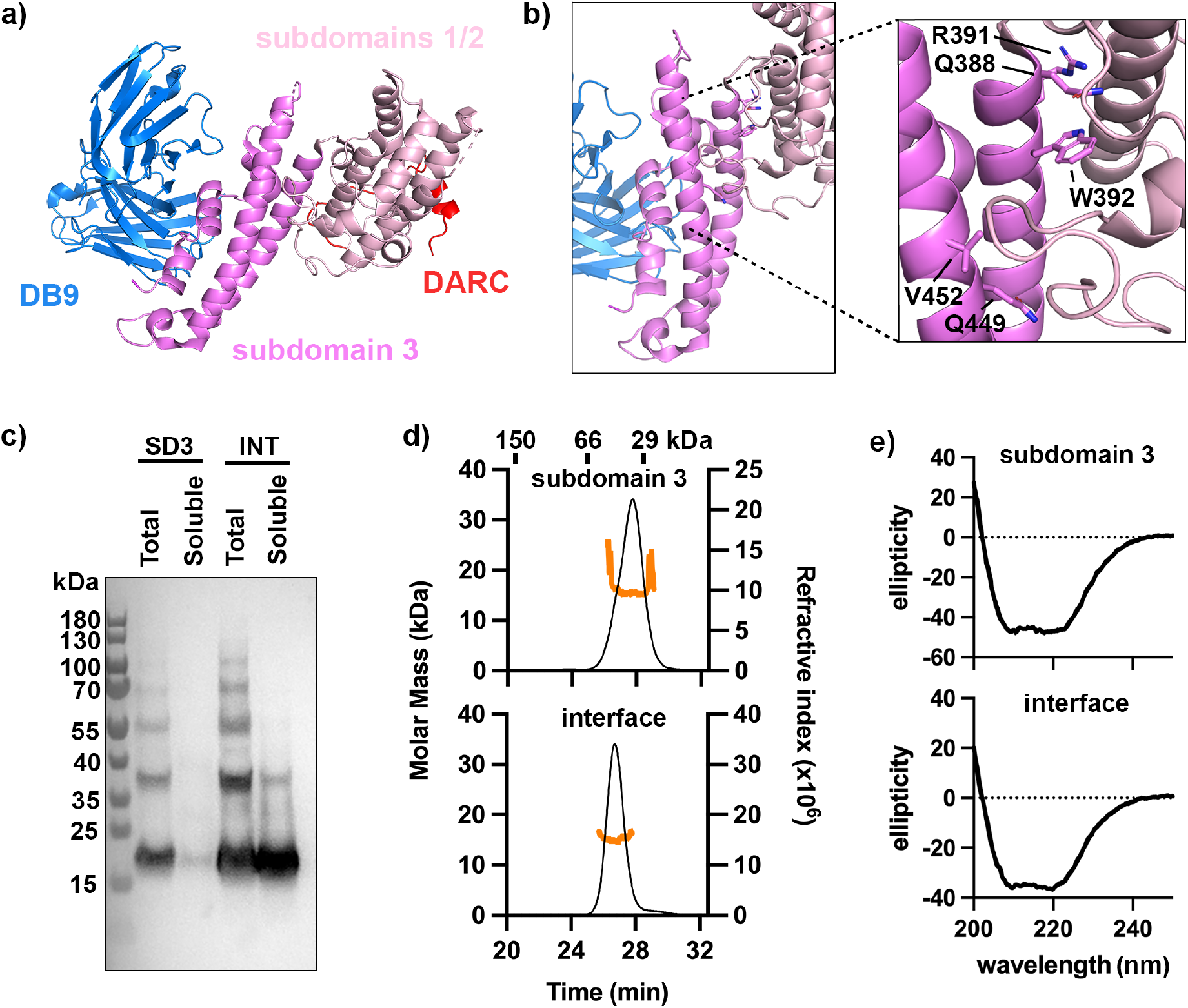
Design of stable variants of PvDBP subdomain 3. **a)** A composite model of PvDBP-RII with subdomain 3 in pink and subdomains 1 and 2 in light pink. A peptide from DARC is shown in red. The variable domains of inhibitory antibody DB9 are shown in blue. This is derived from a composite of PDB codes 6R2S and 8A44, aligned on PvDBP-RII. **b)** A close-up of subdomain 3 of PvDBP-RII, showing the five residues which contact subdomain 2 and are mutated in the interface protein. **c)** A Western blot showing expression of subdomain 3 (SD3) and interface (INT) in *E. coli*. In each case, Total is whole lysed cells while Soluble is whole lysed cells centrifuged with the supernatant loaded onto the blot. **d)** Analysis by SEC-MALS of subdomain 3 and interface. In each case the black line shows the absorbance for the sample analysed on a Superdex 75 column while the orange line shows the mass determined by light scattering. **e)** Analysis by circular dichroism of subdomain 3 and interface proteins showing a predominantly α-helical fold.

Human neutralising monoclonal antibodies can also target epitopes on different subdomains of PvDBP-RII. One study isolated monoclonal antibodies from a human volunteer from a malaria endemic region, finding that these bind predominantly to subdomain 2 and overlap with the binding site for DARC^19-30^ and the proposed dimerization site^18^. In contrast, a second study isolated a panel of ten monoclonal antibodies from human volunteers vaccinated with PvDBP-RII, and showed that one of these, DB9, was most effective at neutralising blood stage growth of a sequence diverse set of *Plasmodium vivax* parasites. DB9 binds to the outer surface of subdomain 3, distant from the characterised DARC binding site^9^. This study also showed that antibodies which target subdomain 2 can antagonise the function of subdomain 3 targeting antibodies, leading to the question of whether vaccination with subdomain 3 alone may be desirable^9^. In this study, we test this hypothesis. We use structural insight to design a protein immunogen which contains just subdomain 3 of PvDBP-RII and assess whether this is a better vaccine immunogen than intact PvDBP-RII.

## Results

### Surface remodelling generates a soluble version of subdomain 3

Subdomain 3 of PvDBP-RII forms an autonomous structural unit, consisting of two long anti-parallel α-helices, along which runs a region of loops and short helices, suggesting that it might be possible to generate a well-expressing version of subdomain 3 which folds correctly (Figure 1a,b). Subdomain 3 packs against subdomain 2 through hydrogen bonds and a small hydrophobic patch. We reasoned that expressing subdomain 3 alone would expose this small hydrophobic patch and might impact its solubility. We therefore designed a version of subdomain 3 in which we resurfaced the hydrophobic patch by replacing hydrophobic residues with hydrophilic alternatives (W392K and V452E) (Figure 1b). In addition, we altered three residues within this interface region to increase their charge (Q388D, R391E and Q449E), with the aim of increasing the solubility of the isolated domain. We name this variant interface and compared it with unaltered subdomain 3.

We expressed both interface and subdomain 3 in *E. coli*. Small scale expression trials demonstrated that subdomain 3 expressed in an insoluble form in the *E. coli* pellet, while interface was expressed in a soluble form (Figure 1c). We next purified both proteins. In the case of interface, we purified the soluble component, while subdomain 3 was refolded from inclusion bodies. This yielded 3 mg from each litre of *E. coli* for subdomain 3 and 20 mg per litre for interface. Both purified proteins were monomeric and monodispersed as demonstrated by SEC-MALLS (Figure 1d) and both showed circular dichroism spectra characteristic of α-helical proteins (Figure 1e), with similar thermal stability and with denaturing transitions at >70°C. (Extended Data Figure 1). Therefore, while both subdomain 3 and interface can be produced from *E. coli* in a folded form, subdomain 3 requires refolding from inclusion bodies. In contrast, surface remodelling allows interface to be produced in a readily scalable form by ensuring that it is expressed as a soluble protein.

### Both interface and subdomain 3 bind to antibody DB9

We next used surface plasmon resonance analysis to assess the binding of interface and subdomain 3 to monoclonal antibody DB9, allowing us to determine whether the epitope is correctly folded. We immobilised DB9 onto the surface of a protein A/G-coated chip and flowed increasing concentrations of PvDBP-RII, subdomain 3 and interface over this chip. All three bound with similar dissociation constants in the nanomolar range, with 2.63 nM for PvDBP-RII, 1.11 nM for subdomain 3 and 2.67 nM for interface (Figure 2a and Extended Data Table 1).

**Figure 2:**
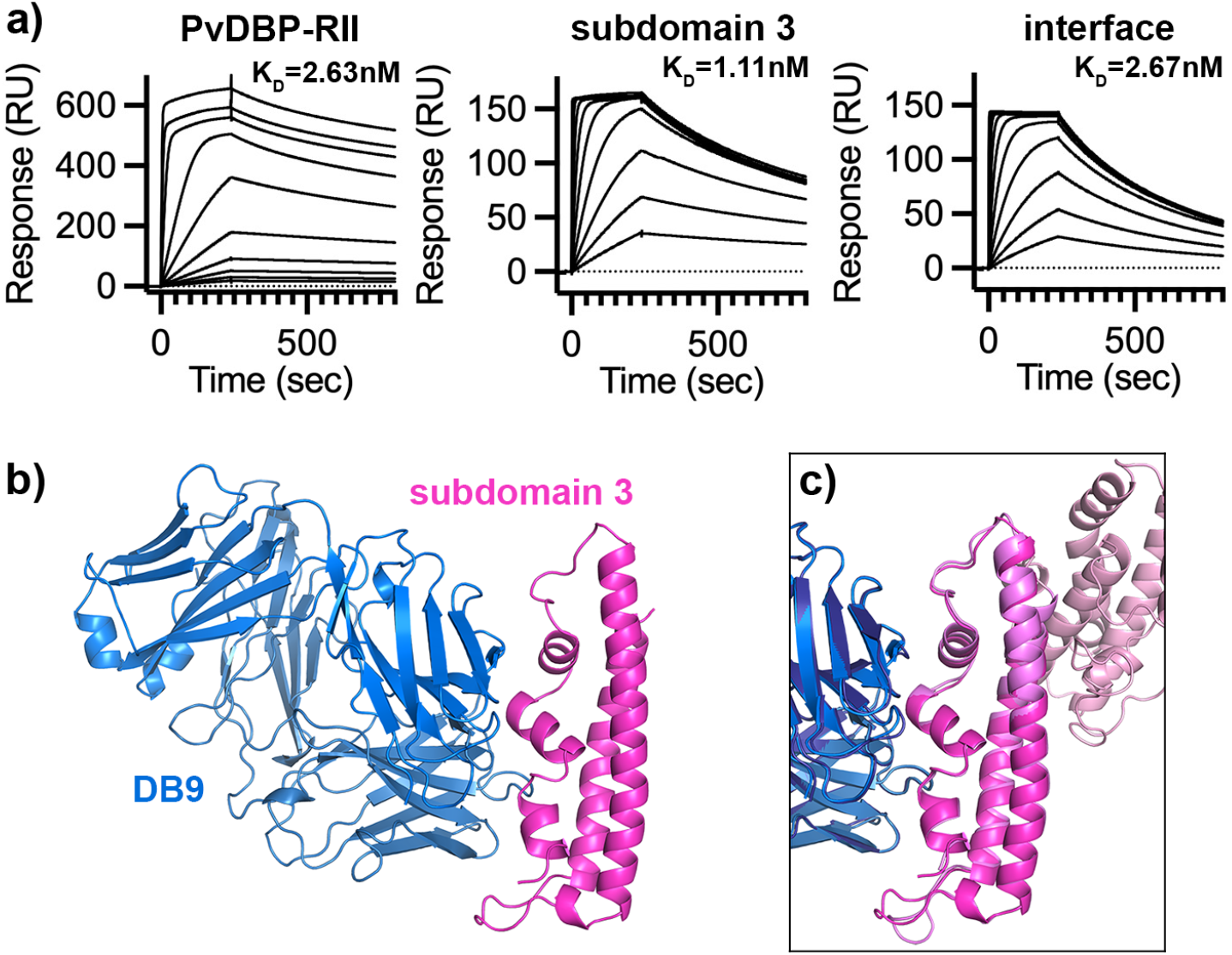
Subdomain 3 and interface bind antibody DB9. **a)** Analysis by surface plasmon resonance of the binding of PvDBP-RII, subdomain 3 and interface to immobilised monoclonal antibody DB9. Each set of curves shows a 2-fold dilution series from a maximum concentration of 1μM. **b)** The structure of subdomain 3 (bright pink) in complex with the Fab fragment of antibody DB9 (blue). **c)** An overlay of the structure of subdomain 3 bound to the Fab fragment of DB9 with that of PvDBP-RII (with subdomain 3 in pink and subdomains 1 and 2 in light pink) bound to the Fab fragment of DB9 (dark blue).

To check the conformation of refolded subdomain 3 we determined its structure using x-ray crystallography (Figure 2b and Extended Data Table 2). We prepared Fab fragments from DB9 and mixed with subdomain 3. Crystals formed and the structure was determined by molecular replacement. Alignment of this structure with that of PvDBP-RII bound to DB9 showed them to align with a root mean square deviation of 0.43Å. A similar approach did not yield crystals of interface, most likely because some of the mutations which generate the interface were involved in forming crystal contacts within the subdomain 3:DB9 crystals.

Both interface and subdomain 3 can therefore be generated in a correctly folded form which retain the ability to bind to neutralising antibody DB9.

### Interface and subdomain 3 generate a more potent neutralising antibody response than PvDBP-RII

We next compared the antibody responses induced in rabbits following immunisation of two rabbits with four 20μg doses of either interface, subdomain 3 or PvDBP-RII. In each case, the immunogens were mixed with Freund’s adjuvant and dosing was conducted on days 0, 14, 28, and 42, with sera harvested on day 56.

To test the efficacy of these sera at preventing erythrocyte invasion, we used a *Plasmodium knowlesi* model. While *Plasmodium vivax* cannot be cultured, transgenic *Plasmodium knowlesi* in which the three PkDBP proteins have been replaced with PvDBP from the SalI strain can be studied using *in vitro* growth inhibitory assays^10^. When used to analyse a panel of monoclonal antibodies, these transgenic lines gave similar outcomes to an *ex vivo Plasmodium vivax* invasion assay^9^.

We started by purifying total IgG from the rabbit sera and used ELISA to assess the quantity of PVDBP-RII and subdomain 3 binding IgG in each sample. IgG from rabbits immunised with interface and subdomain 3 showed similar, or greater ELISA reactivity against both ligands than IgG from rabbits immunised with PvDBP-RII (Extended Data Figure 2). We therefore proceeded to assess their efficacy in a growth inhibition assay, studying two-fold dilutions of total IgG purified from sera from a maximum concentration of 10mg/ml. IgG from rabbits immunised with PvDBP-RII produced only 20-30% growth inhibition at 10mg/ml. In contrast, IgG from rabbits immunised with subdomain 3 and interface were substantially more effective, giving 100% growth inhibition at 10mg/ml and with IC^50^ values of around 5mg/ml (Figure 3a). Therefore subdomain 3 and interface were equivalently effective as vaccine immunogens and both outperformed PvDBP-RII.

**Figure 3:**
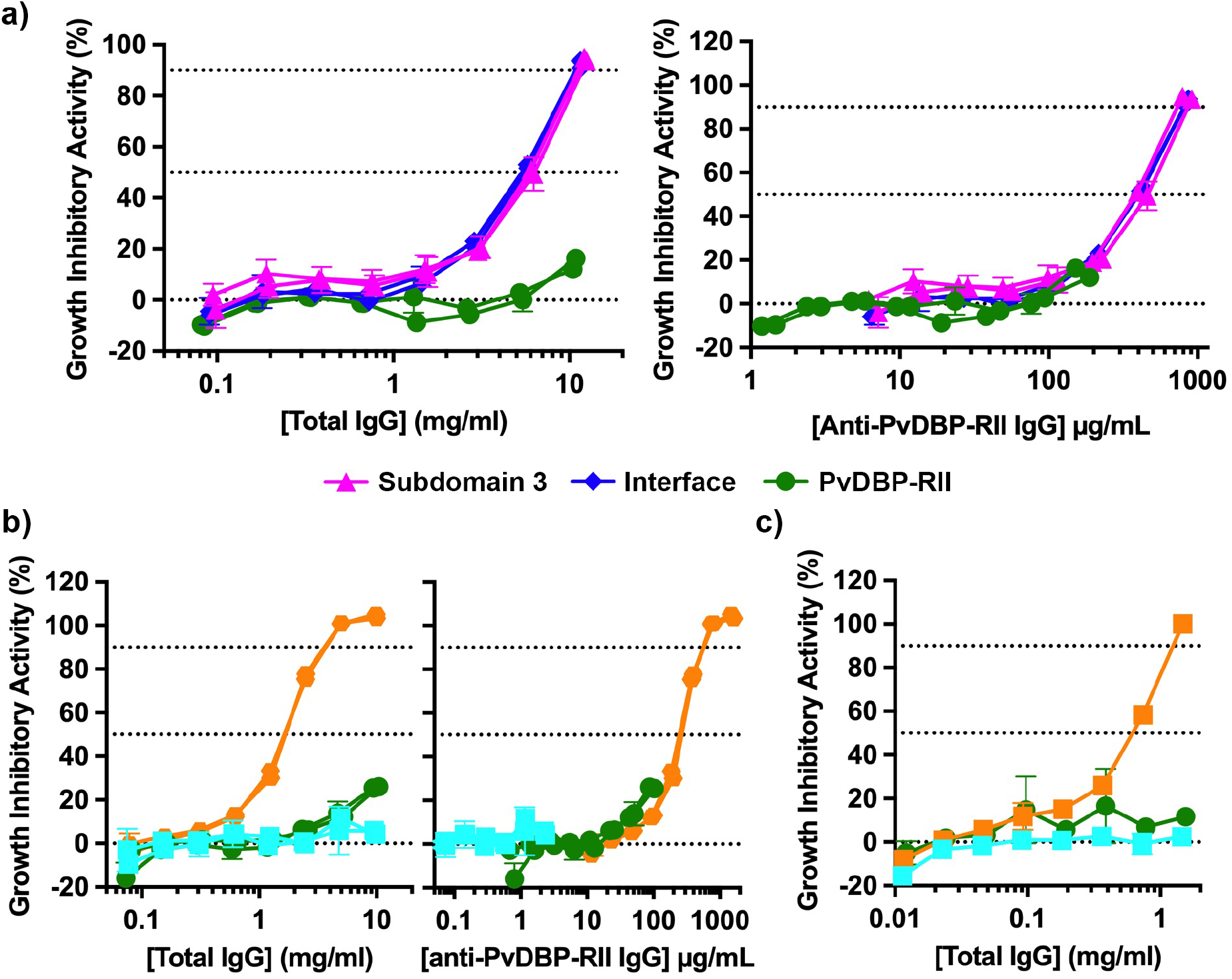
Subdomain 3-based immunogens are more effective than PvDBP-RII. **a)** Growth inhibitory activity for different concentrations of IgG induced in rabbits through immunisation with subdomain 3 (pink), interface (blue) and PvDBP-RII (green), tested against a *Plasmodium knowlesi* strain in which the PkDBPs have been deleted and replaced with PvDBP from the SalI strain of *Plasmodium vivax*. The left show growth inhibitory activity measured against total IgG concentration while the right panel show the same data corrected for the specific quantity of PvDBP-RII specific IgG. Each sample was studied with three technical replicates and these are representative data from n=2. **b)** Total IgG (green), purified from rabbits immunised with PvDBP-RII, was separated using a column displaying subdomain 3 into subdomain 3-specific IgG (orange) and IgG depleted of antibodies binding subdomain 3 (pale blue). Their growth inhibitory activity was measured against *Plasmodium knowlesi* expressing PvDBP from the SalI strain. The left-panel shows the inhibitory activity as a factor of total IgG for each sample, which the right-hand panel is corrected for IgG specific for PvDBP-RII. **c)** An equivalent depletion experiment conducted for a human serum sample from a volunteer vaccinated with PvDBP-RII.

We next asked whether the better performance of subdomain 3 and interface was due to the induction of a larger titre of PvDBP-targeting antibody (i.e. antibody quantity) or due to a greater proportion of growth-inhibitory antibody (i.e. antibody quality). To answer this, we used calibration-free concentration analysis to assess the quantity of PvDBP-RII specific IgG in each sample. This was done using surface plasmon resonance, with PvDBP-RII conjugated to the chip surface at high density to ensure conditions in which mass transport effects were evident, followed by injection of IgG at two different flow rates. Replotting the growth-inhibition data as a factor of PvDBP-RII-specific IgG showed that the curves for rabbits immunised with all three immunogens were equivalent, albeit with PvDBP-RII not inducing sufficient antibodies to reach 50% growth-inhibition (Figure 3a). Therefore, the quality of antibodies induced by PvDBP-RII, interface and subdomain 3 are all equivalent and the improved performance of both interface and subdomain 3 is due to high quantities of growth-inhibitory antigen specific IgG.

The finding that subdomain 3-based immunogens induce IgG of a similar quality to PvDBP-RII led to us to ask whether the growth-inhibitory antibodies induced by PvDBP-RII immunisation all target subdomain 3 or whether there are also growth-inhibitory antibodies targeting other regions of PvDBP-RII. To answer this question, we conducted depletion experiments. Total IgG from rabbits immunised with PvDBP-RII, or from human volunteers immunised with PvDBP-RII as part of a clinical trial, were passed over a column which had been coupled to subdomain 3 protein. The IgG which did not stick to the column (subdomain 3-depleted IgG) and those eluted from the column (subdomain 3-specific IgG) were assessed for the presence of antibodies that bind to subdomain 3 by ELISA, confirming efficient depletion (Extended Data Figure 2). These antibodies were next tested for growth-inhibitory activity, using the transgenic *Plasmodium knowlesi* expressing PvDBP from the SalI variant. Both rabbit (Figure 3b) and human (Figure 3c) subdomain 3-specific IgG showed effective growth inhibition with EC^50^ values of ∼1mg/ml. In contrast, in neither case was growth inhibition observed for subdomain 3 depleted IgG at the maximum concentration that could be achieved. Therefore, all detectable growth-inhibition obtained from either human or rabbit IgG from PvDBP-RII-immunised individuals was due to antibodies binding to subdomain 3.

## Conclusions

Controlled human malaria infection of volunteers vaccinated with a protein vaccine consisting of the PvDBP-RII immunogen formulated with Matrix M adjuvant have provided the first evidence that PvDBP-based vaccines can affect the multiplication rate of *Plasmodium vivax* in vaccinated humans^19^. Nevertheless, this study also highlights that current vaccines fall short of inducing the levels of immunity required for sterile protection and emphasises the need for improved PvDBP-based vaccine immunogens^19^. Here, we attempt to produce such an immunogen using a rational, structure-guided approach.

Structure-guided vaccine design often starts with structural studies to reveal how the most effective neutralising, or in this case growth-inhibitory, monoclonal antibodies function. In the case of PvDBP-RII, studies have been conducted of both mouse and human antibodies, resulting in structures of the epitopes of five antibodies^9,18,21,25^. These studies are much smaller in scope those conducted for other malaria antigens, such as PfRH5 and PfCSP, where hundreds of monoclonal antibodies have been analysed, and the outcomes are less clear. Antibodies that bind to various regions of PvDBP can be growth-inhibitory, including those that target subdomain 2^18^, where the DARC binding site^21^ and proposed dimerization interface lie^22^, or those that target subdomain 3^9^. It is also not clear how each of these growth-inhibitory antibodies functions, with steric hinderance of membrane approach proposed as a possible mechanism for those that target subdomain 3^9^. Despite this, we decided to follow up our finding that the broadly-reactive, growth-inhibitory antibody DB9 binds to subdomain 3 of PvDBP-RII^9^ and to design and test subdomain 3 as a vaccine immunogen.

Subdomain 3 adopts a discrete three α-helical architecture which interacts with other parts of PvDBP-RII through a small hydrophobic patch. Subdomain 3 alone expresses in an insoluble form in bacteria and required refolding to produce a functional protein. In contrast, resurfacing of the exposed hydrophobic patch, through five amino acid changes, resulted in a soluble, stable, highly expressed subdomain 3 immunogen, which we call interface. In our hands, PvDBP-RII is challenging to express and is often not stable on storage, which may be limiting its effectiveness as an immunogen after formulation with adjuvant or immunisation of human volunteers. In contrast, interface is extremely stable and scalable production is likely to be effective.

Side-by-side comparison in a well-established model of growth-inhibition revealed interface to induce more effective growth-inhibitory responses than PvDBP-RII. Indeed, at the maximum concentration of IgG tested, PvDBP-RII was 20-30% effective and interface was 100% effective, with an equivalent EC^30^ around five-fold lower for interface. When we separated antibodies induced using PvDBP-RII into those that did and did not bind to subdomain 3, we found that growth-inhibitory antibodies predominantly bound to subdomain 3. Indeed, when we assessed the growth-inhibitory effect of antibodies induced using PvDBP-RII and interface, as a function of subdomain 3-reactive antibodies, we found that PvDBP-RII and interface performed equivalently. Therefore, to our surprise, the better efficacy of antibodies induced by subdomain 3 was not due to improvement in the quality of the antibody response, through focusing onto a more effective epitope region. Instead, both PvDBP-RII and interface induced predominantly subdomain 3-targeting growth-inhibitory antibodies and interface induced these in greater quantities.

In summary, this study uses rational, structure-based immunogen design to produce a novel form of PvDBP which is stable, readily produced and induces a more growth-inhibitory antibody response than previous PvDBP-based immunogens. This is now available for clinical testing as a component of vaccines to prevent *Plasmodium vivax*.

## Materials and methods

### Expression and purification of PvDBP-RII, subdomain 3 and interface

PvDBP-RII and antibody DB9 were produced as described previously^9^. Gene sequences for PvDBP-RII, subdomain 3 and interface were obtained from GeneArt and were cloned into a modified version of the pET15b vector to provide an N-terminal his-tag followed by a TEV cleavage site. These were expressed in *E. coli* BL21-DE3 cells, induced with 1mM IPTG at OD 1.0 and grown overnight at 18°C.

Subdomain 3 was found in the insoluble fraction and was purified by refolding. Cells were resuspended in 20mM Tris pH 8.0, 300mM NaCl, 20mM imidazole and broken by sonication. After centrifugation at 50,000g for 30 minutes, the pellet was resuspended in 6M GdnHCl, 20mM Tris pH 8, 20mM imidazole, 10mM β-mercaptoethanol by incubation at room temperature for 2 hours before centrifugation at 50,000g for 30 minutes at 4°C. The soluble fraction was incubated with Ni-NTA beads, and the bound material was washed in the 6M GdnHCl, 20mM Tris pH 8, 20mM imidazole, 300μM oxidised glutathione, 3mM reduced glutathione. It was then refolded while attached to the Ni-NTA column with a slow decreasing concentration of GdnHCl, while maintaining other buffer components unchanged, before eluting in 20mM Tris pH 8, 300 mM NaCl, 200mM imidazole. This yielded ∼3mg of protein per litre of cells.

The sequence of interface is: PDIYEKIREWGRDYVSELPTEVQKLKEKCDGKIAYTDKK VCKVPPCQNACKSYDQWITRKKNEWDELSNKFISVKNAEKVQTAGIVTPYDILKQELDEFNEVAFENEINKRDGAYIELCVC. Interface was expressed in a soluble form. Cells were lysed as for subdomain 3 and the soluble fraction was applied to a Ni-NTA column. This was washed using 20mM Tris pH 8, 300mM NaCl, 20mM imidazole and the protein was eluted using 20mM Tris pH 8, 300 mM NaCl, 200mM imidazole, yielding around 20mg per litre of cells.

All proteins were next dialysed into PBS and cleaved with TEV protease overnight at room temperature. They were then passed through a Ni-NTA column and the flow through was collected. This was concentrated and applied to a Superdex 75 (Cytiva) in 20mM Tris pH 8.0, 150mM NaCl.

### Circular dichroism analysis

Circular dichroism analysis was conducted using a J-815 spectropolarimeter (JASCO, Japan) with an attached Peltier water bath. Proteins were buffer exchanged using PD10 columns into 10mM sodium phosphate pH 7.5, 150mM NaF and were diluted to a final concentration of 0.2mg/ml. Spectra were collected from 190nm to 260nm wavelengths at 25°C and four independent measurements were averaged together to obtain the final curve. To study thermal stability, spectra were collected from 200nm and 250nm wavelengths at 2°C intervals at temperatures from 20°C to 90°C. The ellipticity at 220nm wavelength was plotted against temperate to determine the melting temperature.

### Surface plasmon resonance

Surface plasmon resonance analysis was conducted using a Biacore T200 instrument (GE healthcare) using 20mM HEPES pH 7.4, 150mM NaCl, 0.005% Tween-20. A Biacore chip was prepared by using amine coupling to coat a CM5 chip in protein A/G. Monoclonal antibody DB9 was then captured onto flow path 2, with flow path 1 left as a negative control. To analyse binding of PvDBP-RII, subdomain 3 and interface to DB9, these were then flowed across the chip surface using a two-fold dilution series from a maximum concentration of 1μM. Data were analysed using the BIAevaluation software.

### Crystallisation and structure determination

For crystallisation, subdomain 3 and the Fab fragment of DB9 were mixed in a ratio of 1.1:1 and were incubated at room temperature for 30 minutes. The mixture was loaded onto a superdex 200 10/30 column, run in 20mM Tris pH 8.0, 50mM NaCl (Cytiva). The protein was concentrated to 11mg/ml and subjected to crystallisation trials. Crystals grew with reservoir solution of 0.2M ammonium acetate, 0.1M sodium acetate pH 4.0, 15% PEG 4000. These were transferred into a cryo-protection solution containing 0.2M ammonium acetate, 0.1M sodium acetate pH 4.0, 15% PEG 4000, 25% glycerol. A dataset was collected to a final resolution of 1.55Å on beamline I03 at Diamond Light Source. Molecular replacement was conducted in Phaser^26^ using the previous structure of the Fab fragment of DB9 bound to PvDBP-RII (PDB:6R2S)^9^ as a search model, after trimming to leave the DB9 Fab fragment bound to subdomain 3 with loops removed. Modelling building was conducted in coot^27^ and refinement in buster^28^.

### Immunisation of rabbits

Before immunization, all protein constructs underwent endotoxin removal using Pierce High-capacity endotoxin removal resin (ThermoFisher Scientific). Subsequently, the constructs were dispatched to GenScript (Oxford, UK) for rabbit immunization. Intramuscular immunization was performed on two rabbits with 20 µg of PvDBP-RII, subdomain 3 or interface, each administered with Freund’s adjuvant. The dosing regimen consisted of four doses administered at two-week intervals. Serum samples were collected from all rabbits two weeks after the final dose.

### Measurement of binding by ELISA

Qualitative IgG binding ELISAs were carried out by coating PvDBP-RII or subdomain 3 on Maxisorp flat-bottom 96-well ELISA plates (Nunc) at 2 µg/mL in 50 µL at 4 °C overnight. Plates were then washed twice with PBS and 0.05% Tween 20 (PBS/T) and blocked with 200 µL of Blocker™ Casein (Thermo Fischer Scientific) for 1 h. Next, wells were incubated with 10 µg/mL of sera for approximately 1 hr at 20 °C then washed 4 times with PBS/T before the addition of 50 µL of 1:1000 dilution of goat anti-rabbit gamma-chain alkaline phosphatase-conjugated secondary antibody (Sigma-Aldrich) for 1 hr at 20 °C. Wells were then washed 6 times with PBS/T and developed with 100 µL of p-nitrophenyl phosphate substrate at 1 mg/mL (Sigma-Aldrich) and optical density read at 405 nm (OD_405_) using a Model 550 Microplate Reader (Bio-Rad, UK).

### Assays of growth inhibitory activity using Plasmodium knowlesi lines

#### In vitro parasite culture and synchronisation

Human RBC-adapted parasites were maintained in culture as previously described^29^. Briefly, parasites were grown at 2 % haematocrit in O+ human RBC, which were prepared twice monthly. Culture medium contained 10 % heat-inactivated pooled human serum mixed with RPMI 1640 supplemented with 25 mM HEPES, 35 μM hypoxanthine, 2 mM L-glutamine and 20 μg/mL gentamycin. Parasite cultures were synchronised at trophozoite/schizont stage by magnetic separation (MACS LS columns, Miltenyi Biotech).

#### In vitro assay of Growth Inhibitory Activity

The assay was adapted from the protocol of the International GIA Reference Centre at NIH, USA^30^. Synchronised trophozoites were adjusted to 1.5 % parasitaemia, and 20 μl aliquots were pipetted into 96-well flat/half area tissue culture cluster plates (Appleton Woods). 20 μl test antibody or controls were added in duplicate or triplicate test wells over a concentration range (usually; 1, 0.5, 0.25, 0.125, 0.0625, 0.0312, 0.015 and 0.0075 mg/ml) and incubated for one erythrocytic parasite cycle (26-30 h). Parasitaemia was measured using the lactate dehydrogenase (pLDH) activity assay following standard protocols^31^. Percentage GIA was calculated as below;

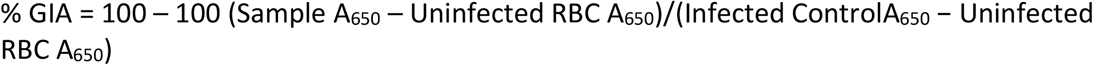

An anti-DARC VHH camelid nanobody^32^, a kind gift from Dr Olivier Bertrand (INSERM, France) was included in the test plate as a positive control in every assay (at a final concentration of 6, 3 or 1.5 μg/mL) and an anti-*Ebolavirus* glycoprotein-reactive human IgG1 mAb as a negative isotype control.

The human serum sample came from a healthy UK adult vaccinated with a PvDBP-RII protein/adjuvant in the VAC079 clinical trial. The trial was conducted at the University of Oxford and received ethical approval from UK National Health Service Research Ethics Services and regulatory approval from the UK Medicines and Healthcare products Regulatory Agency^19^.

#### Calibration-free concentration analysis (CFCA)

Calibration-free concentration analysis (CFCA) was conducted using a Biacore T200 instrument with a Biacore Biotin CAPture sensor chip (Cytiva) following established procedures^33^. Prior to experimentation, all antigen and total IgG samples were buffer exchanged into HBS EP+ running buffer (10 mM HEPES pH 7.4, 150 mM NaCl, 0.3 mM EDTA, and 0.05% Tween-20) utilizing Zeba Spin desalting columns (ThermoFisher Scientific). Biotin CAPture reagent was immobilized on the chip at a flow rate of 2 µl/min for 120 s on two flow paths. Subsequently, 10 µg/ml of biotinylated PvDBP-RII was captured on the sensor at a flow rate of 2 µl/min for 600 s, ensuring > 2000 response units (RU) of antigen on the flow cell. These parameters were optimized to saturate the sensor chip completely, thus achieving mass transport effects. Following antigen capture, a stabilization period of 600 s was maintained at the same flow rate. For flow path 2, 1 mg/ml of total IgG sample, the concentration determined using absorbance at 280 nm, was passed over the antigen-coated sensor at a flow rate of 5 µl/min for 36 s to measure the initial rate of antigen-specific binding, while flow path 1 served as a negative control. Chip regeneration was performed using the manufacturer’s supplied regeneration reagent (diluted 1:2 in HBS EP+). Each sample was analyzed twice, once with a total IgG sample flow rate of 5 µl/min and again at 100 µl/min, as CFCA analysis requires measuring initial rates of antigen-specific binding at slow and fast flow rates.

Data analysis was conducted using the Biacore T200 analysis software with the CFCA analysis function. Measurements with initial binding rates between 0.3 – 15 RU/s and QC (quality control) values > 0.13, indicating sufficient mass transport limitation, were selected, following the recommendations of the manufacturer. The binding model assumed a molecular weight of 150 kDa for IgG, and the diffusion coefficient of IgG at 20 °C in HBS EP+ was set at 4.8 x 10^-11^ m2/s, as previously determined^33^.

## Acknowledgements

This work was funded by a Wellcome Investigator Award (20797/Z/20/Z) to MKH. NMB was funded by the Wellcome-funded PhD program in Cellular Structural Biology. TP was funded by a Skaggs-Oxford graduate scholarship. The authors would like to thank David Staunton for support with biophysics data collection and Dr Ed Lowe and the beamline scientists at Diamond I03 for support with crystallographic data collection. We thank Rob Moon for the *Plasmodium knowlesi* strains. We also thank Mimi Hou, Angela Minassian and Chetan Chitnis for provision of the VAC079 clinical trial serum.

## Data availability

Data within graphs (source data) and uncropped gel and blot images are included with this submission. The crystal structure is deposited in the Protein Data Bank with an accession code of 9EZE.

---

**Extended Data Figure 1:**
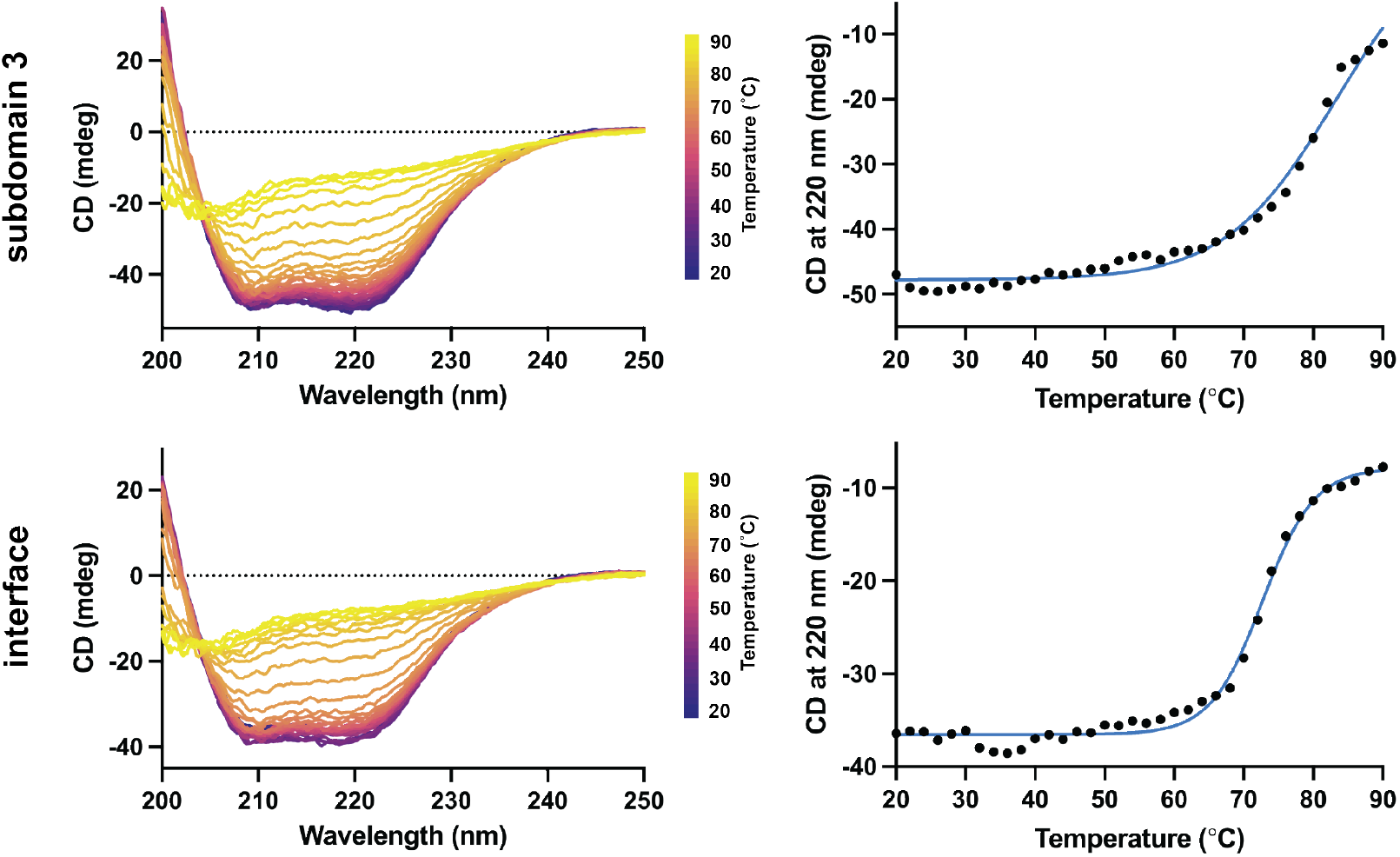
Thermal melts of subdomain and interface. Left: the thermal melt curves between wavelengths 200 and 250 nm, at temperatures from 20°C to 90°C. Right: The thermal denaturation of the alpha-helical protein was measured at 220 nm to estimate the melting temperature (T_m_).

**Extended Data Figure 2:**
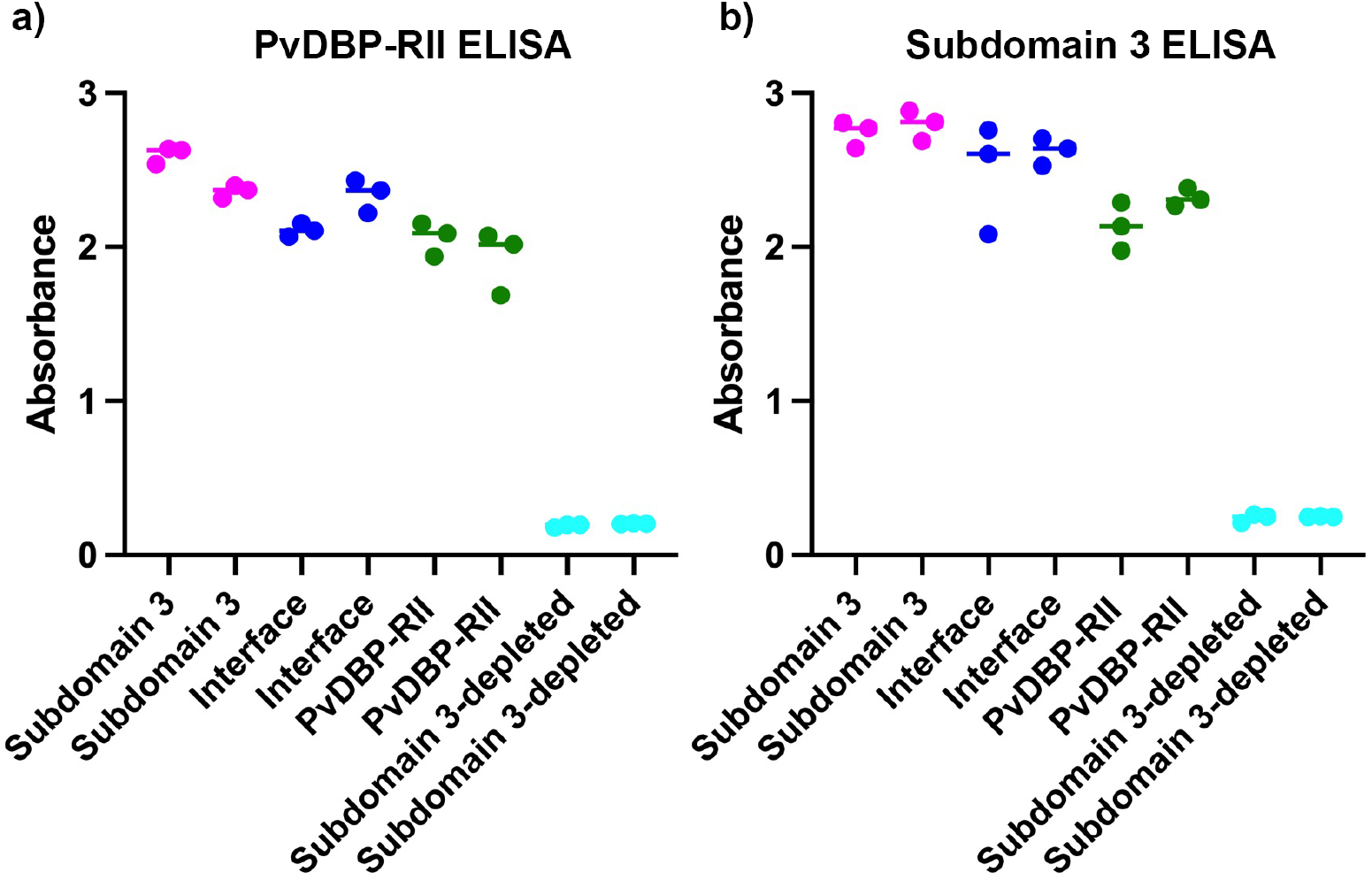
ELISA measurements for purified IgG. ELISA was used to measure the binding of IgG from immunised rabbits against immobilised**a)** PvDBP-RII and **b)** Subdomain 3. In each case, we studied two rabbits immunised with Subdomain 3, with interface or with PvDBP. Each sample was studied with three technical replicates. We also depleted subdomain 3-binding antibodies from the PvDBP-RII IgG, yielding the subdomain 3-depleted IgG.

**Extended Data Table 1:**
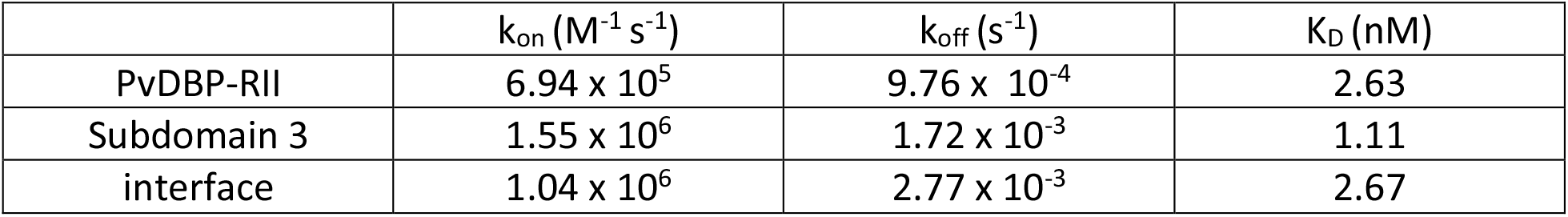
kinetic parameters measured by surface plasmon resonance analysis for binding to monoclonal antibody DB9

**Extended Data 2:** crystallographic statistics

|  |  |
| --- | --- |
| **Data collection** |  |
| Space group | P2<sub>1</sub>2<sub>1</sub>2<sub>1</sub> |
| Cell dimensions: |  |
| a, b, c (Å) | 76.43, 126.19, 129.86 |
| α, β, γ (°) | 90.00, 90.00, 90.00 |
| Resolution (Å) | 90.50 – 1.55 (1.605-1.55) |
| Total observations | 1178345 (112071) |
| Total unique | 180370 (17671) |
| R<sub>pim</sub> | 2.36 (30.77) |
| CC<sub>1/2</sub> | 0.999 (0.58) |
| I/σ(I) | 14.77 (1.96) |
| Completeness (%) | 98.30 (95.67) |
| Multiplicity | 6.5 (6.3) |
| Wilson B factor (Å<sup>2</sup>) | 21.4 |
| **Refinement** |  |
| Number of reflections | 178875 |
| R<sub>work</sub> / R<sub>free</sub> | 20.4 / 22.3 |
| Average B factor (Å<sup>2</sup>) | 30.0 |
| Number of residues: |  |
| Amino acid residues | 1107 |
| Waters | 978 |
| RMSZ deviations |  |
| Bond lengths | 0.011 |
| Bond angles | 1.12 |
| Ramachandran plot |  |
| Favoured (%) | 98.1 |
| Allowed (%) | 1.9 |
| Outliers (%) | 0.0 |

## References

1. Battle, K.E. et al. Mapping the global endemicity and clinical burden of Plasmodium vivax, 2000-17: a spatial and temporal modelling study. Lancet 394, 332–343 (2019).

2. Howes, R.E. et al. Plasmodium vivax Transmission in Africa. PLoS Negl Trop Dis 9, e0004222 (2015).

3. Price, R.N., Douglas, N.M. & Anstey, N.M. New developments in Plasmodium vivax malaria: severe disease and the rise of chloroquine resistance. Curr Opin Infect Dis 22, 430–5 (2009).

4. Flannery, E.L., Markus, M.B. & Vaughan, A.M. Plasmodium vivax. Trends Parasitol 35, 583–584 (2019).

5. De, S.L., Ntumngia, F.B., Nicholas, J. & Adams, J.H. Progress towards the development of a P. vivax vaccine. Expert Rev Vaccines 20, 97–112 (2021).

6. Horuk, R. et al. A receptor for the malarial parasite Plasmodium vivax: the erythrocyte chemokine receptor. Science 261, 1182–4 (1993).

7. Miller, L.H., Mason, S.J., Clyde, D.F. & McGinniss, M.H. The resistance factor to Plasmodium vivax in blacks. The Duffy-blood-group genotype, FyFy. N Engl J Med 295, 302–4 (1976).

8. Singh, A.P. et al. Targeted deletion of Plasmodium knowlesi Duffy binding protein confirms its role in junction formation during invasion. Mol Microbiol 55, 1925–34 (2005).

9. Rawlinson, T.A. et al. Structural basis for inhibition of Plasmodium vivax invasion by a broadly neutralizing vaccine-induced human antibody. Nat Microbiol 4, 1497–1507 (2019).

10. Mohring, F. et al. Rapid and iterative genome editing in the malaria parasite Plasmodium knowlesi provides new tools for P. vivax research. Elife 8(2019).

11. Chitnis, C.E. & Sharma, A. Targeting the Plasmodium vivax Duffy-binding protein. Trends Parasitol 24, 29–34 (2008).

12. Chitnis, C.E. & Miller, L.H. Identification of the erythrocyte binding domains of Plasmodium vivax and Plasmodium knowlesi proteins involved in erythrocyte invasion. J Exp Med 180, 497–506 (1994).

13. de Cassan, S.C. et al. Preclinical Assessment of Viral Vectored and Protein Vaccines Targeting the Duffy-Binding Protein Region II of Plasmodium Vivax. Front Immunol 6, 348 (2015).

14. Moreno, A. et al. Preclinical assessment of the receptor-binding domain of Plasmodium vivax Duffy-binding protein as a vaccine candidate in rhesus macaques. Vaccine 26, 4338–44 (2008).

15. Cole-Tobian, J.L. et al. Strain-specific duffy binding protein antibodies correlate with protection against infection with homologous compared to heterologous plasmodium vivax strains in Papua New Guinean children. Infect Immun 77, 4009–17 (2009).

16. King, C.L. et al. Naturally acquired Duffy-binding protein-specific binding inhibitory antibodies confer protection from blood-stage Plasmodium vivax infection. Proc Natl Acad Sci U S A 105, 8363–8 (2008).

17. Nicolete, V.C., Frischmann, S., Barbosa, S., King, C.L. & Ferreira, M.U. Naturally Acquired Binding-Inhibitory Antibodies to Plasmodium vivax Duffy Binding Protein and Clinical Immunity to Malaria in Rural Amazonians. J Infect Dis 214, 1539–1546 (2016).

18. Urusova, D. et al. Structural basis for neutralization of Plasmodium vivax by naturally acquired human antibodies that target DBP. Nat Microbiol 4, 1486–1496 (2019).

19. Hou, M.M. et al. Vaccination with Plasmodium vivax Duffy-binding protein inhibits parasite growth during controlled human malaria infection. Sci Transl Med 15, eadf1782 (2023).

20. Singh, S.K., Hora, R., Belrhali, H., Chitnis, C.E. & Sharma, A. Structural basis for Duffy recognition by the malaria parasite Duffy-binding-like domain. Nature 439, 741–4 (2006).

21. Moskovitz, R. et al. Structural basis for DARC binding in reticulocyte invasion by Plasmodium vivax. Nat Commun 14, 3637 (2023).

22. Batchelor, J.D., Zahm, J.A. & Tolia, N.H. Dimerization of Plasmodium vivax DBP is induced upon receptor binding and drives recognition of DARC. Nat Struct Mol Biol 18, 908–14 (2011).

23. Batchelor, J.D. et al. Red blood cell invasion by Plasmodium vivax: structural basis for DBP engagement of DARC. PLoS Pathog 10, e1003869 (2014).

24. Chootong, P. et al. Mapping epitopes of the Plasmodium vivax Duffy binding protein with naturally acquired inhibitory antibodies. Infect Immun 78, 1089–95 (2010).

25. Chen, E. et al. Broadly neutralizing epitopes in the Plasmodium vivax vaccine candidate Duffy Binding Protein. Proc Natl Acad Sci U S A 113, 6277–82 (2016).

26. McCoy, A.J. et al. Phaser crystallographic software. J Appl Crystallogr 40, 658–674 (2007).

27. Emsley, P., Lohkamp, B., Scott, W.G. & Cowtan, K. Features and development of Coot. Acta Crystallogr D Biol Crystallogr 66, 486–501 (2010).

28. Bricogne G. B.E., Brandl M., Flensburg C., Keller P. P. W. and Roversi P, Sharff A., Smart O.S., Vonrhein C. ‘BUSTER version 2.10.4’. Cambridge, United Kingdom: Global Phasing Ltd. (2017).

29. Moon, R.W. et al. Adaptation of the genetically tractable malaria pathogen Plasmodium knowlesi to continuous culture in human erythrocytes. Proc Natl Acad Sci U S A 110, 531–6 (2013).

30. Miura, K. et al. Anti-apical-membrane-antigen-1 antibody is more effective than anti-42-kilodalton-merozoite-surface-protein-1 antibody in inhibiting plasmodium falciparum growth, as determined by the in vitro growth inhibition assay. Clin Vaccine Immunol 16, 963–8 (2009).

31. Kennedy, M.C. et al. In vitro studies with recombinant Plasmodium falciparum apical membrane antigen 1 (AMA1): production and activity of an AMA1 vaccine and generation of a multiallelic response. Infect Immun 70, 6948–60 (2002).

32. Smolarek, D. et al. A recombinant dromedary antibody fragment (VHH or nanobody) directed against human Duffy antigen receptor for chemokines. Cell Mol Life Sci 67, 3371–87 (2010).

33. Payne, R.O. et al. Human vaccination against Plasmodium vivax Duffy-binding protein induces strain-transcending antibodies. JCI Insight 2(2017).

